# Infection of point-of-lay hens to assess the sequential events during H7N7 high-pathogenicity avian influenza emergence at a layer premises

**DOI:** 10.1101/2025.07.16.665166

**Authors:** Amanda H. Seekings, Elizabeth Billington, Sahar Mahmood, Saumya S. Thomas, Anita Puranik, Simon Johnson, Alejandro Nunez, Samantha Watson, Jill Banks, Sharon M. Brookes, Ian H. Brown, Marek J. Slomka

## Abstract

H7N7 low-pathogenicity avian influenza virus (H7N7-LPAIV) incursions have preceded emergence of H7N7 high-pathogenicity (HP)AIV at several European layer hen outbreaks. Evidence from a UK layer H7N7-HPAIV (2008) outbreak informed in vivo modelling of sequential events, beginning with H7N7-LPAIV precursor incursion. Three groups of 17-week-old (point-of-lay) hens were inoculated with three precursor H7N7-LPAIV candidates which possessed: Group 1 – the classic LPAIV single-basic cleavage site (CS) (H7N7-SBCS); Group 2 - the dibasic CS (H7N7-DBCS) discovered at the outbreak (both represented direct H7N7-HPAIV (2008) precursors); Group 3 - a related European H7N7-LPAIV (2008) which possessed a DBCS within its haemagglutinin (HA) gene which was highly-conserved with the HA of the UK (2008) outbreak H7N7-HPAIV, but its other genetic segments differed. In Groups 1-3, initial H7N7-LPAIV exposure caused limited viral shedding, restricted H7-antibody responses and no deaths, by 14-days post-inoculation (dpi). H7N7-HPAIV challenge of all groups (n=8 per challenge group, including AIV-naïve hens in Group 4) followed at 14 dpi. Prior H7N7-LPAIV inoculation protected against mortality due to H7N7-HPAIV challenge to varying degrees, with eight (100%), seven (87.5%), six (75%) and three (37.5%) survivors in Groups 1-4, respectively, by study-end (14-days post-challenge), when all 24 survivors had strongly seroconverted. Among challenge survivors belonging to Groups 1-3, 20/21 (95%) shed H7N7-HP, indicating that immunity acquired during initial H7N7-LPAIV exposure did not arrest H7N7-HPAIV replication. Despite prior H7N7-LPAIV incursion which elicited additional clinical protection against H7N7-HPAIV challenge, we showed how an emergent H7N7-HPAIV may represent a source for onward spread and further HPAIV outbreaks.

## 1. Introduction

Avian influenza viruses (AIVs) are influenza A viruses of avian origin, containing eight single-stranded negative-sense RNA genetic segments (Knipe and Howley, 2013). AIVs are subtyped antigenically according to the surface glycoproteins haemagglutinin (16 HA subtypes) and neuraminidase (nine NA subtypes) (Alexander, 2007). Low-pathogenicity (LP)AIVs circulate in their natural wild bird reservoir, with no overt clinical signs. LPAIV poultry incursions cause infection which is self-resolved and normally restricted to the respiratory and enteric tracts, with mild or inapparent clinical outcomes (Swayne and Halvorson, 2008). The H5 and H7 AIV subtypes can potentially mutate in galliforme hosts from LPAIV to the corresponding high-pathogenicity (HP)AIVs, where outbreaks in commercial galliformes (such as chickens) cause high morbidity and mortality due to systemic viral dissemination, with accompanying economic damage to the poultry industry (Alexander and Brown, 2009).

The LPAIV to HPAIV change may occur in H5 and H7 subtypes, via genetic mutation within the HA gene’s cleavage site (CS). Likely molecular mechanisms causing this mutation have been reviewed (de Bruin et al., 2022), where HA CS extension is produced by insertion of several basic amino acids into the LPAIV single-basic (SB)CS to produce a HPAIV multi-basic (MB)CS (OFFLU, 2022). The HA glycoprotein precursor, HA0, requires post-translational host-protease cleavage to be functional and produce infectious viral progeny (Garten, 2018; Rott, 1992; Steinhauer, 1999). The LPAIV SBCS is cleaved by trypsin-like proteases which are restricted to the avian respiratory and enteric tracts, generally limiting replication to these tissues. However, MBCS generation makes the HA susceptible to cleavage by ubiquitous furin-like proteases, causing extensive HPAIV systemic spread and virulent pathogenesis in galliformes (Garten and Klenk, 2008).

H5Nx HPAIVs of the “goose/Guangdong” (GsGd) lineage (Brown et al., 2006; World Health Organization Global Influenza Program Surveillance, 2005) remain the major cause of HPAI outbreaks worldwide, where the current clade 2.3.4.4b H5N1 HPAIV remains a major global concern (Caliendo et al., 2022; Nunez and Ross, 2019; Pan American Health Organization, 2023), with epizootics affecting whole countries for up to several months (Byrne et al., 2023;

Pohlmann et al., 2022). However, the GsGd H5N1 prototype strain first emerged approximately 30 years ago, but no closely related or precursor LPAIV was identified (Wan, 2012). H7-HPAIV outbreaks are also economically damaging, but within the last 20 years these have been largely restricted in well-resourced countries, with statutory responses limiting outbreaks to relatively few (or even single) premises (Abdelwhab et al., 2014). Interestingly, European H7N7-HP outbreaks since 2008 have provided clear evidence for initial H7N7-LPAIV incursion and intra-flock circulation, followed by mutation to generate the homologous H7-HPAIV within the same (Bonfanti et al., 2014; Byrne et al., 2021; Mulatti et al., 2017; Seekings et al., 2018) or a nearby epidemiologically linked premises (Dietze et al., 2018). All these European H7N7-HPAIV outbreaks included index cases at outdoor layer premises, which may provide important clues to ultimately unravel the circumstances under which the LPAIV to HPAIV switch may occur. Retrospective examination of flock morbidity, including an earlier drop in egg-laying at the UK (2008) H7N7-HPAIV outbreak, indicated initial H7N7-LPAIV incursion approximately two weeks before suspect HPAI clinical signs led to H7N7-HPAIV confirmation (Seekings et al., 2018). H7-specific serology at these European H7N7-HPAIV outbreaks strongly supported an epidemiological sequence of events, where H7N7-LPAIV spread among layers, with minimal clinical signs, leading to seroconversion and LPAI resolution. Analyses of the four epidemiological units at the UK (2008) outbreak indicated that H7N7-HPAIV emergence and spread occurred against a background of differing H7-specific seroprevalences in the four units, whereby prior H7 immunity ameliorated HPAI clinical signs (Seekings et al., 2018).

An important discovery during the UK (2008) outbreak was the presence of H7N7-LPAIV sequences, originating from environmental (faecal deep litter) samples, exclusively consisting of a di-basic (DB)CS (Seekings et al., 2018). The DBCS has been rarely encountered among H7 LPAIVs and contains two successive K residues at the HA CS, immediately prior to the conserved RGLF motif. The DBCS is classified as a LPAIV, so it may represent an intermediate between the more frequently observed SBCS among H7 LPAIVs, and the corresponding emergent H7 HPAIV (OFFLU, 2022). Despite extensive investigation at the UK H7N7-HPAIV outbreak, no SBCS H7N7-LPAIV precursor was discovered (Seekings et al., 2018). Interestingly, the DBCS occurred in an H7N7-LPAIV wild mallard strain, also isolated during 2008 (Sweden), where the HA gene shared strong (98.5%) nucleotide similarity, although the other seven viral genetic segments were distinct (Metreveli et al., 2010; Seekings et al., 2018).

Our current study aimed to model the sequential events at the UK (2008) layer outbreak, beginning with H7N7-LPAIV incursion, followed 2-weeks later by H7N7-HPAIV challenge. Unlike other HPAIV in vivo investigations which typically utilise younger chickens (James et al., 2023; Seekings et al., 2024; Seekings et al., 2023; Seekings et al., 2021), one important detail was to inoculate older point-of-lay hens, to reflect the bird characteristics of the UK (2008) H7N7-HPAIV layer outbreak. Epidemiological findings from this outbreak informed our novel study design, where reverse genetics (RG) provided two candidate LPAIV precursors of the outbreak HPAIV, one with the discovered DBCS and the other with a SBCS. The SBCS, DBCS and MBCS (HPAIV) viruses were otherwise genetically identical (including the internal genes) (Seekings et al., 2020), and used to compare the impact of precursor LPAIVs with different CSs on subsequent HPAIV challenge. We also used the DBCS Swedish (2008) isolate as a third H7N7-LPAIV, to compare the effects of prior LPAIV exposure to a virus with differing internal genes. Virological and pathogenesis outcomes were monitored throughout the initial H7N7-LPAIV and subsequent H7N7-HPAIV phases.

## 2. Materials and methods

### 2.1 Ethics and biosafety

Animal experiments were approved by the Animal and Plant Health Agency (APHA) Animal Welfare and Ethical Review Body, in accordance with UK legislation. Experiments were carried out in UK approved SAPO level 4, ACDP level 3 biocontainment laboratories and animal facilities at APHA-Weybridge, UK, including engineering controls with strict adherence to safe working practices and use of personal protective equipment (PPE) by animal caretakers. To respect animal welfare, monitoring for humane endpoints was done twice daily following H7N7-HPAIV challenge, and increased to thrice daily following severe disease onset, thereby informing euthanasia decisions to reduce unnecessary suffering and instances of “found dead” hens (Seekings et al., 2023). For any found dead hens, the death time was recorded as having occurred mid-way since the preceding welfare inspection. Hens were housed on straw bedding, with access to food and water ad libitum, with the latter changed daily.

### 2.2 Viruses

H7N7 HPAIV A/chicken/England/11406/2008 (abbreviated as H7N7-HPAIV; GISAID accession numbers EPI712884-EPI712891) was isolated during the UK (2008) layer outbreak (Seekings et al., 2018).

Three H7N7-LPAIVs were used. The native H7N7 LPAIV A/mallard/Sweden/100993/2008 (mall/SE/08; GISAID accession numbers EPI1232143-EPI1232150) was selected because it contained the same DBCS as the LPAIV precursor to the UK (2008) H7N7-HPAIV layer outbreak (Metreveli et al., 2010; Seekings et al., 2018). Reverse genetics (RG) rescued two viable recombinant LPAIVs derived from the H7N7-HPAIV. Both RG viruses were genetically identical to the isolated H7N7-HPAIV except for the CS, with one containing a DBCS (H7N7-DBCS) and the other a SBCS (H7N7-SBCS) (Seekings et al., 2020). Viral stocks were grown in 9-day-old specified pathogen free (SPF) embryonated fowls’ eggs (Valo, Germany) (WOAH, 2021)), the allantoic fluids harvested and sterile filtered (0.2µm, Sartorius, Germany), with viral titres determined as a 50% egg infectious dose (EID50) (Reed and Muench, 1938). The stocks were further diluted in sterile 0.1M phosphate-buffered saline, to provide inocula for the in vivo investigations, and also the respective antigens for the haemagglutination inhibition assay.

### 2.3 Experimental design

Fifty-two point-of-lay Novogen Brown hens (Tom Barron Ltd., Preston, UK) of high health status were pre-screened for AIV exposure prior to inoculation by collecting swabs (buccal and cloacal), for testing by M-gene RRT-PCR (below). Sera were also pre-screened from wing bleeds, for testing by a commercial ELISA, which detects anti-influenza type A-specific antibodies to the viral nucleoprotein (NP) (IDEXX, France, as per the manufacturer’s instructions). These tests served to exclude active or previous AIV infection in the hens.

To model the sequential events observed during the H7N7-HPAIV UK (2008) layer outbreak (Seekings et al., 2018), for the initial H7N7-LPAIV phase, three groups of hens (n=14 per group, at 17-weeks-age) modelled H7N7-LPAIV incursion into layers. The three groups were segregated and housed in separate pens throughout the H7N7-LPAIV phase, which began with concurrent inoculation with the three H7N7-LPAIVs, H7N7-SBCS (Group 1), H7N7-DBCS (Group 2) or mall/SE/08 (Group 3; Figure 1). The inoculum for each hen contained 1 × 106 EID50 of the respective H7N7-LPAIV in 100µl, administered via the intranasal / intraocular route. The Group 1-3 hens were swabbed (buccal and cloacal) daily, until 12 days post-inoculation (dpi). Rigorous discipline was strictly applied during daily husbandry and sampling by animal caretakers throughout the entire study, including cleaning and replacement of external PPE upon completion of these tasks in one pen, before proceeding to the next. To investigate any H7N7-LPAIV pathogenesis, two hens from each of Groups 1-3 were randomly selected for cull at 2 and 4 dpi for tissue sampling. At the end of the H7N7-LPAIV phase, at 14 dpi, the ten remaining hens in Groups 1-3 were again swabbed and were wing bled for serological testing.

**Fig. 1.**
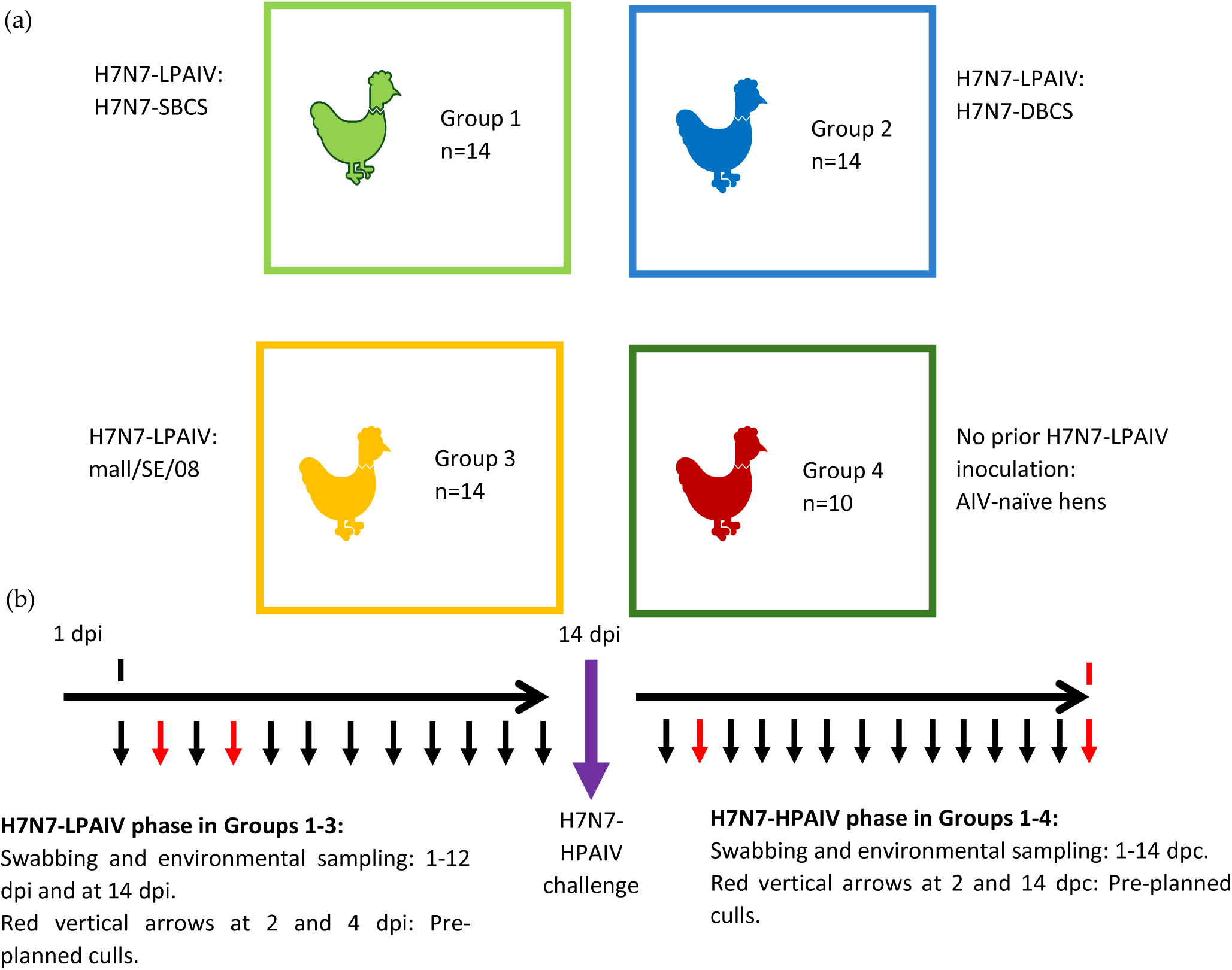
Schematic diagram of the in vivo study design. (a) the four groups of hens, where 14 hens were inoculated with the indicated H7N7-LPAIV in Groups 1-3. (b) The study timeline indicates daily swabbing and environmental sampling during the initial H7N7-LPAIV phase (indicated by vertical arrows). The thicker vertical purple arrow at 14 days post-infection (dpi) indicates swabbing and wing bleeds at the end of H7N7-LPAIV phase in Groups 1-3, followed immediately on the same day by H7N7-HPAIV challenge of Groups 1-4. The subsequent H7N7-HPAIV phase included daily monitoring and swabbing until study-end at 14 days post-challenge (dpc; corresponding to 28 dpi for Groups 1-3). Red vertical arrows on the timeline during the H7N7-LPAIV phase (Groups 1-3) indicate pre-planned culls of two hens per group on each day at 2 and 4 dpi, and during the H7N7-HPAIV phase (Groups 1-4) at 2 dpc (two apparently healthy hens per group) and 14 dpc (two surviving hens per group, at study-end). The culls enabled organ sampling, with bloods collected as indicated.

To address the effects of the subsequent H7N7-HPAIV emergence during the UK (2008) layer outbreak (Seekings et al., 2018), the ten hens in Groups 1-3 were challenged with H7N7-HPAIV (1 × 106 EID50 per hen, as above) at 14 dpi (corresponding to 0 days post-challenge (dpc); Figure 1). Group 4 served as the age-matched AIV naïve control group (n=10), and had been housed in a separate containment room, away from Groups 1-3, during the preceding H7N7-LPAIV phase. The four groups of hens therefore remained strictly segregated throughout the entire study, with stringent biosecurity protocols maintained across the successive H7N7-LPAIV and H7N7-HPAIV phases, to avoid risk of unintended virus transfer between groups. Daily swabbing followed until study-end at 14 dpc, corresponding to 28 dpi for Groups 1-3, relative to previous inoculation with the three H7N7-LPAIVs. Two hens which displayed no overt clinical signs at 2 dpc were humanely culled for H7N7-HPAIV pathogenesis investigation (tissue sampling), leaving up to eight hens to be monitored in each group until study-end, when two hens per group were humanely culled for tissue sampling. Tissues were also obtained from mortalities following H7N7-HPAIV challenge. Survival of hens within the four groups during the H7N7-HPAIV phase was compared by the Kaplan-Meier method, using the log-rank test, by using PRISM software (GraphPad, CA, USA). All survivors were heart bled at study-end, for serological testing.

### 2.4 Clinical scoring

All live hens were assessed daily for semi-quantitative (subjective) scoring of their health status, with scoring recorded up to twice daily following H7N7-HPAIV challenge. Absence of any overt clinical signs scored as 3 for each hen; mild signs (any combination of ruffled feathers, lethargy, huddling or dropped wings) scored as 2; and moderate signs (mild signs plus any combination of cyanosis, closed eyes or diarrhoea) registered 1. Severe signs scored as 0, typified by stronger neurologic manifestations which prompted euthanasia, including tremors, torticollis and / or an inability to source food and water autonomously. When scoring individual hens, only the most severe score was recorded per day, and enabled calculation of the average clinical health score for all individuals in each group:.

### 2.5 Processing clinical and environmental samples

Swabs (buccal and cloacal) were collected daily and expressed into 1ml brain-heart infusion broth containing antibiotics (BHIB) (Slomka et al., 2018). Portions of 17 internal organs (trachea, nasal turbinates, lung, duodenum, liver, pancreas, colon, caecum, caecal tonsil, bursa, spleen, thymus, oviduct, heart, brain, kidney and breast muscle) were excised at post-mortem, cut and similarly stored at 10% (v/v) in 1ml BHIB, including feather calami (Slomka et al., 2012). Total RNA was extracted robotically from swabs and tissues (Slomka et al., 2009). Except for the caecum and colon, portions of the same tissues together with skin, ovary, air sacs, jejunum, proventriculus and gizzard samples were also collected at post-mortem and fixed in 10% (v/v) buffered formalin.

During both the H7N7-LPAIV and H7N7-HPAIV phases, two environmental samples were collected daily from each group, these being drinking water from a bell drinker (replaced daily) and a mix of faeces and straw. The solid matrices were suspended at 10% (v/v) in BHIB, with centrifugation at 1000g for 5 minutes followed by supernatant removal for robotic total RNA extraction (Slomka et al., 2009). Drinking water was robotically extracted without BHIB addition.

### 2.6 Identification of infected hens using matrix (M)-gene reverse transcription RealTime (RRT)-PCR testing

To detect H7N7 viral RNA (vRNA), total RNA extracts from swabs were tested by the matrix (M)-gene RRT-PCR (Spackman et al., 2002), as described (Slomka et al., 2009). H7N7 vRNA was quantified as relative equivalent units (REUs) by using a 10-fold dilution series of extracted vRNA from EID50-quantified allantoic fluid of H7N7-HPAIV, to provide a standard curve (Londt et al., 2008). Negative swab results were “No Ct” (Ct 40) by M-gene RRT-PCR, while positive viral shedding included swab Ct values ≤36 (Seekings et al., 2018; Slomka et al., 2010; Slomka et al., 2009). Sub-threshold Ct values (36.01–39.99) were included as evidence of shedding (supplementary information, (Slomka et al., 2018)). Any instance of viral shedding from either tract (buccal or cloacal) classified an individual hen as infected. To assess the overall H7N7-HPAIV shedding following challenge, the mean vRNA levels detected from both tracts in each group were compared by area under the curve (AUC) analysis (Seekings et al., 2024).

### 2.7 Tissue tropism

M-gene RRT-PCR testing was also applied to total RNA extracted from tissues. Formalin-fixed tissue sections were examined for histopathological changes and for influenza A virus antigen by virus-specific immunohistochemical (IHC) investigation (Londt et al., 2008).

### 2.8 Serology

Sera derived from clotted bloods were incubated at 56°C for 30 minutes and tested by haemagglutination inhibition (HI) using four haemagglutination units (WOAH, 2021). At the end of the H7N7-LPAIV phase (14 dpi), sera from Groups 1-3 were tested by the respective homologous H7N7-LPAIV antigen. Evidence of H7N7-LPAIV infection included homologous H7-HI reciprocal titres of 16 (or greater) interpreted as positive, but also any instances of H7 reactivity (partial seroconversion, <1:16) were noted at this acute stage, as per similar serological criteria used previously (Slomka et al., 2018). Sera collected at study-end (14 dpc (28 dpi)) from H7N7-HPAIV challenge survivors were H7-HI tested, using the homologous H7N7-HPAIV antigen.

## 3. Results

### 3.1 Inoculation of hens with H7N7-LPAIVs

Evidence obtained during the UK 2008 H7N7-HPAIV layer outbreak guided in vivo modelling of sequential events, which occurred during H7N7-HPAIV emergence. Three potential H7N7-LPAIV precursors were initially inoculated into layers, divided into three groups (Groups 1, 2 and 3; n=14 hens per group; Figure 1), using H7N7-SBCS, H7N7-DBCS and mall/SE/08, respectively. Two hens from each group were culled at 2 and 4 dpi (H7N7-LPAIV phase), and at 2 dpc (H7N7-HPAIV phase), for pathogenesis investigations, leaving eight hens per group for monitoring through both phases of the study. M-gene RRT-PCR testing of swabs identified infection in one (12.5%), two (25%) and five (62.5%) of eight hens in each of Groups 1, 2 and 3 respectively, during the H7N7-LPAIV phase (Figure 2 and Table 1). The remaining 16 hens in Groups 1-3 were considered uninfected by any of the three H7N7-LPAIVs, based on the absence of any shedding from either tract, along with the absence of any serological reactivity by H7-HI testing of sera collected at 14 dpi (Figure 3 and Table 1), but were noted as having been H7N7-LPAIV exposed. All hens remained apparently healthy throughout the H7N7-LPAIV phase.

**Fig. 2.**
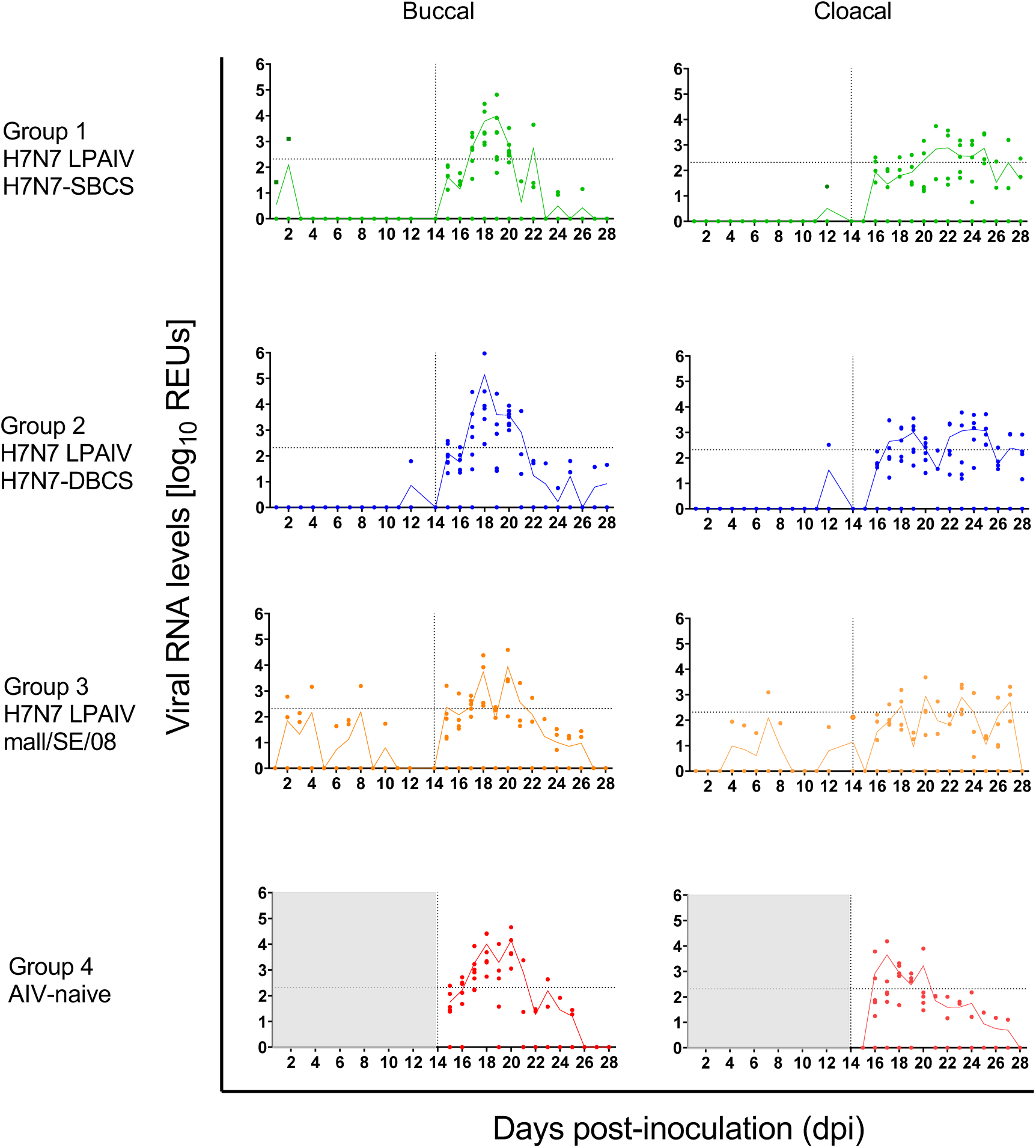
Viral shedding (buccal and cloacal) from hens during the initial H7N7-LPAIV and subsequent H7N7-HPAIV phases. Viral shedding was determined by M-gene RRT-PCR testing and measured as REUs. H7N7-LPAIV shedding was monitored in ten hens until 14 dpi in Groups 1-3, following inoculation with H7N7-SBCS, H7N7-DBCS and mall/SE/08, respectively. Grey shaded area indicates that no swabs were taken from hens in Group 4 during 1-14 dpi. H7N7-HPAIV challenge of all four groups followed at 14 dpi (vertical dotted line), with viral shedding monitored daily until study-end at 14 dpc (28 dpi for Groups 1-3) for the eight hens per group, including AIV naïve hens (Group 4). Shedding levels are shown for individual hens by symbols. Mean shedding levels are indicated by continuous lines, for all live hens on a given day. Horizontal dotted lines denote the REU equivalent of Ct 36, where lower REU shedding levels were sub-threshold.

**Fig. 3.**
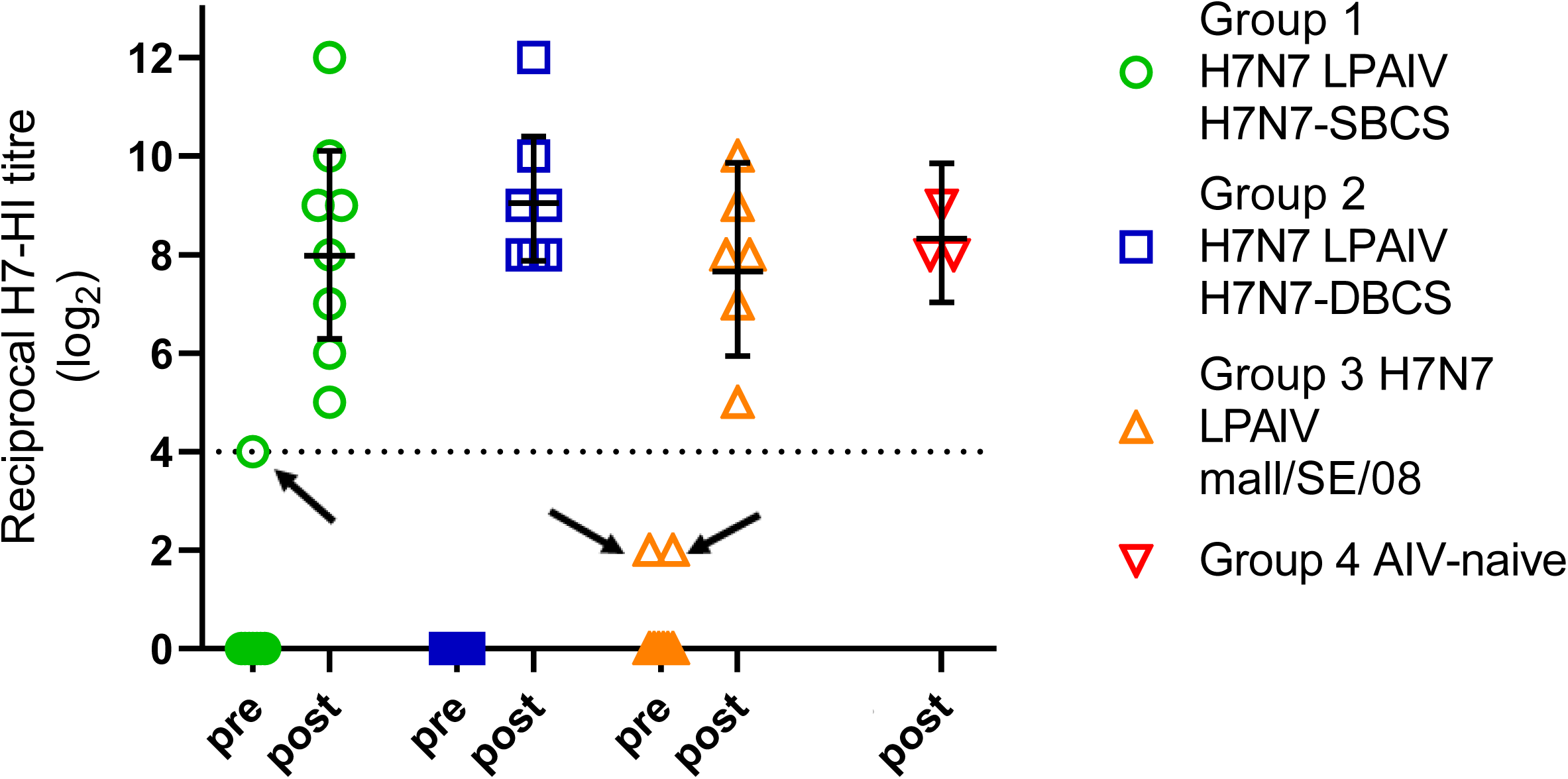
H7-specific antibodies measured at the pre- and post-challenge (H7N7-HPAIV inoculation) time points. Homologous H7-HI titres are shown by symbols in individual hens. Sera were collected from all LPAIV exposed groups at 14 dpi (end of H7N7-LPAIV phase for Groups 1-3) immediately prior to H7N7-HPAIV challenge on the same day, indicated by “pre”. H7-HI titres were also determined at study-end (indicated by “post”, i.e. 14 dpc,) from all surviving hens in the four groups. Homologous H7N7-LPAIV antigens were used for H7-HI testing of sera at 14 dpi from the corresponding LPAIV group (Groups 1-3), while H7N7-HPAIV served as the antigen to test sera collected at 14 dpc (Groups 1-4). Geometric means of the H7-HI titres are shown for sera from each of the groups, along with the 95% confidence interval. The Group 4 hens had been earlier shown to be AIV naïve, see Materials and Methods 2.3. The dotted horizontal line indicates the 1:16 positive H7-HI cut-off, but weaker H7-HI titres (H7 reactors) are included. Arrows indicate three H7-HI reactor sera in Groups 1 and 3 collected at 14 dpi, from hens which shed (Ct<36) during the H7N7-LPAIV phase.

**Table 1.**
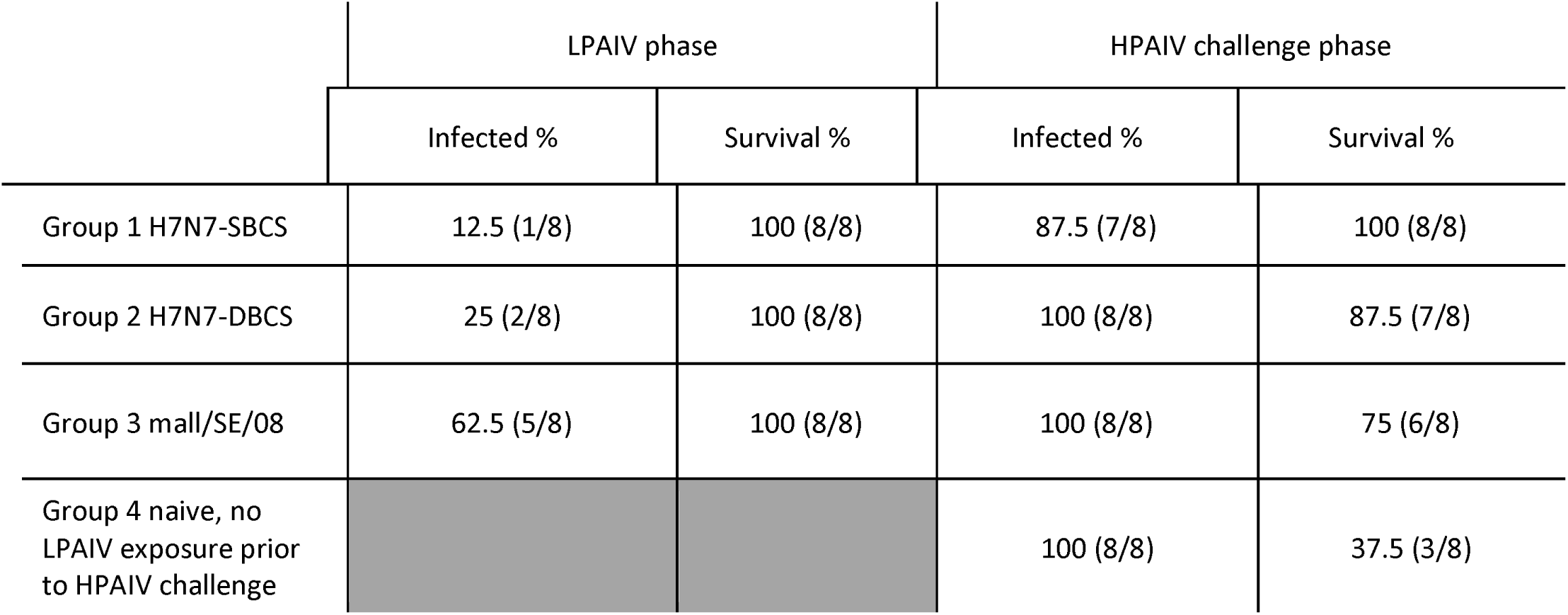
Summary of the infection status and survival of the hens during (i) the H7N7-LPAIV phase of the study (Groups 1-3; 1-14 dpi) and (ii) following H7N7-HPAIV challenge at 14 dpi, which also included Group 4. Infection and survival outcomes for the eight hens per group, which were monitored throughout the entire study period. Infected hens were identified by viral shedding, which included subthreshold Ct values.

|  | LPAIV phase |  | HPAIV challenge phase |  |
| --- | --- | --- | --- | --- |
|  | Infected % | Survival % | Infected % | Survival % |
| Group 1 H7N7-SBCS | 12.5 (1/8) | 100 (8/8) | 87.5 (7/8) | 100 (8/8) |
| Group 2 H7N7-DBCS | 25 (2/8) | 100 (8/8) | 100 (8/8) | 87.5 (7/8) |
| Group 3 mall/SE/08 | 62.5 (5/8) | 100 (8/8) | 100 (8/8) | 75 (6/8) |
| Group 4 naive, no LPAIV exposure prior to HPAIV challenge |  |  | 100 (8/8) | 37.5 (3/8) |

Serology confirmed H7N7-LPAIV infection in one Group 1 hen (#3508; HI titre of 1:16) and partial seroconversion in two Group 3 hens (#3537 and #3559, each HI titre of 1:4) by 14 dpi (Figure 3). No serological reactivity occurred among any of the Group 2 hens. H7N7-LPAIV shedding in Groups 1-3 was largely sporadic at overall low levels. By including sub-threshold shedding, only one hen (#3539, Group 3) demonstrated sustained H7N7-LPAIV shedding for more than two successive days, spanning 3-8 dpi inclusive (Figure 2), but remained H7 seronegative by 14 dpi.

Planned culls of two hens per H7N7-LPAIV inoculated group at 2 and 4 dpi provided 18 organs from each hen, and all were vRNA negative (data not shown). Together with the largely sporadic and low H7N7-LPAIV shedding levels in Groups 1-3, this observation underlined highly limited and localised replication of the three H7N7-LPAIVs, with no detectable vRNA in sampled tissues, including the respiratory and enteric tracts. No histopathological changes were detected, with no virus-specific staining detected by IHC investigation.

### 3.2 H7N7-HPAIV challenge

H7N7-HPAIV challenge investigated the next phase of the UK (2008) outbreak, modelling H7N7-HPAIV emergence within the layers. H7N7-HPAIV was administered at 14 dpi to the previously H7N7 LPAIV-inoculated hens (Groups 1-3), alongside challenge of Group 4 (AIV naïve) hens (Figure 1). Of 10 challenged hens per group, two were culled for tissue sampling at 2 dpc, leaving eight birds to be monitored to study-end (14 dpc). H7N7-HPAIV challenge caused mortalities among hens previously exposed to H7N7-LPAIV in Groups 2 (one death: #3521 at 3.4 dpc, 12.5%) and 3 (two deaths: #3540 and #3538 at 2.4 and 2.8 dpc, respectively, 25%), but no mortalities (0%) in Group 1 (Figure 4, Table 1). These observations demonstrated varying degrees of protection against HPAI mortality among hens previously exposed to the three H7N7-LPAIVs. H7N7-HPAIV mortalities were highest in the previously naïve hens (Group 4), where all eight became infected and five deaths (62.5%) occurred, between 2.4-6.0 dpc. A significant difference in mortality frequency was determined between Groups 1 and 4 (p=0.0085), but the differences between mortality frequencies in Groups 2 and 3 compared to Group 4 were not significant (p>0.05).

**Fig. 4.**
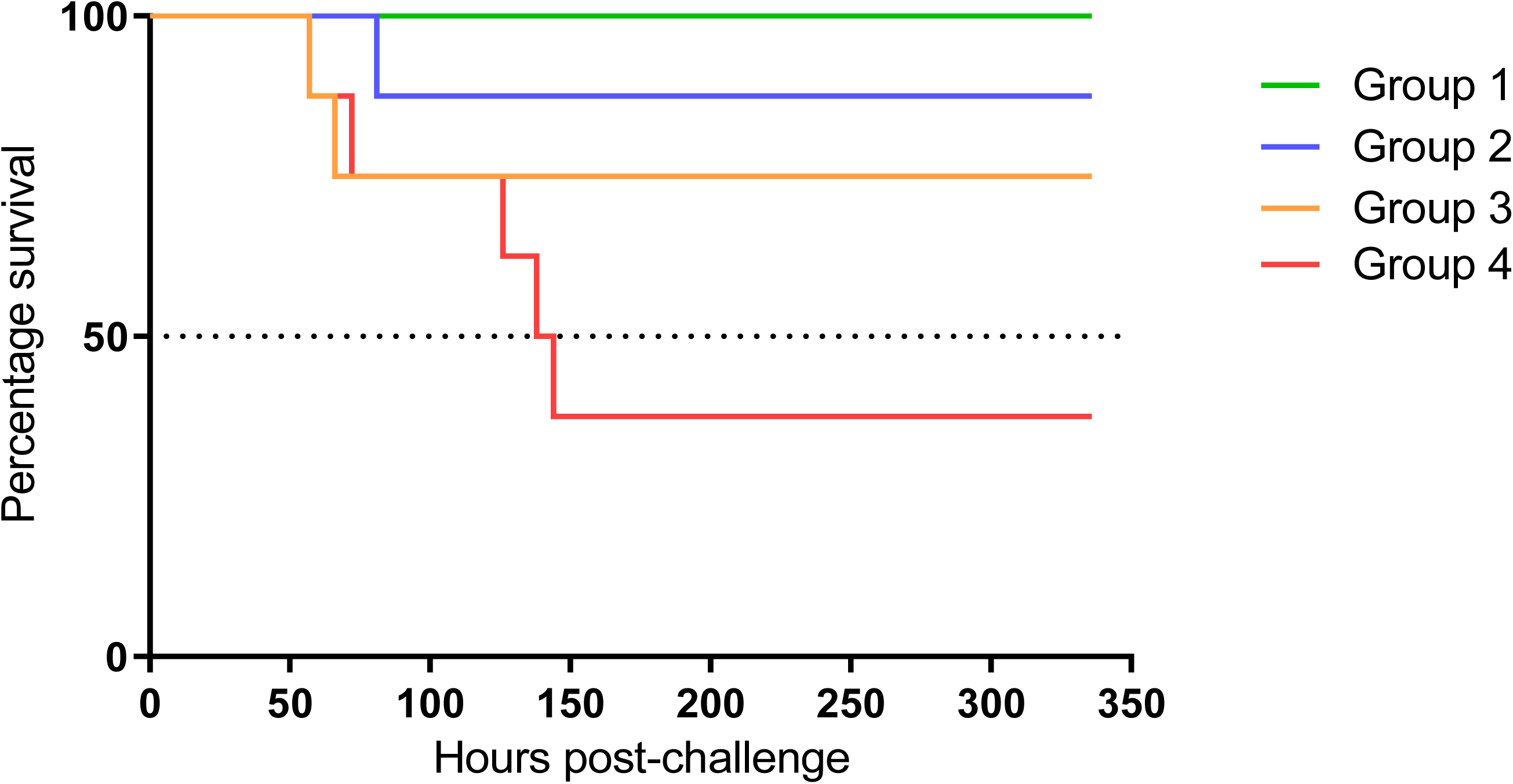
Survival of hens following H7N7-HPAIV challenge. The percentage of surviving hens in Groups 1-4 is shown for the four groups, during the HPAIV phase, where eight hens per group were monitored to study-end at 14 dpc. The two apparently healthy challenged hens per group which were culled at 2 dpc for tissue sampling are not included. For visual guidance, 50% survival is indicated by the dotted horizontal line.

H7N7-HPAIV infection occurred in Groups 1-4, with positive buccal shedding occurring between 1-9 dpc (Figure 2). Positive cloacal shedding was first detected at 2 dpc, and continued until 6 dpc (Group 4), but for Groups 1-3, cloacal shedding endured longer until 14 dpc (Figure 2), suggesting a temporal shift in H7N7-HPAIV shedding tropism from the respiratory to the enteric tract. The five Group 4 mortalities had a mean death time of 4.5 days (Figure 4), where the three survivors successfully resolved shedding and seroconverted by study-end (Figures 2 and 3). Within all four groups, H7N7-HPAIV buccal shedding attained higher peak levels than cloacal shedding, but AUC analyses showed no significant differences between buccal and cloacal shedding (p>0.05).

From shedding data, seven hens in Group 1 (87.5%) were infected with H7N7-HPAIV, with no mortalities (Figure 2, Table 1), and included one H7N7-HPAIV uninfected hen (#3508) which previously shed H7N7-SBCS and had seroconverted to HI titre 1:16 by 14 dpi (Figure 3). In Group 2, all eight hens shed H7N7-HPAIV, while Group 3 also experienced 100% H7N7-HPAIV infection (Table 1). Although all eight Group 3 hens shed H7N7-HPAIV, two survivors experienced only sub-threshold buccal shedding, with both (#3539, #3559) having previously shed mall/SE/08 during the LPAIV phase.

In summary, prior H7N7-LPAIV exposure did not prevent subsequent H7N7-HPAIV infection in 23/24 (95.8%) challenged hens. Twenty-one of the pre-exposed hens (87.5%) survived H7N7-HPAIV challenge and all seroconverted by 14 dpc (Figure 3), including all eight hens in Groups 1-3 that had been classified as previously H7N7-LPAIV infected (Table 1).

### 3.3 Clinical scoring after H7N7-HPAIV challenge

In addition to the H7N7-HPAIV mortalities, the average clinical scores were monitored in the four groups until study-end (Figure 5). All Group 1 hens survived H7N7-HPAIV challenge and displayed only mild signs. The single Group 2 mortality was preceded by moderate signs 12 hours earlier. Clinical signs among the seven Group 2 survivors were mostly mild, although transient moderate signs occurred for under one day in two hens, which then improved clinically. Group 3 hens experienced two (25%) mortalities which occurred suddenly or were preceded 12 hours earlier by mild signs. The six survivors experienced the least clinical signs among the four groups, with no signs apparent between 8 dpc and study-end (Figure 5). The Group 4 (previously AIV naïve) hens experienced not only the greatest mortality (62.5%), but the three challenge survivors displayed more clinical signs compared to the survivors in Groups 1-3, which endured for 0.5-2 days as moderate signs, before improvement to mild signs. There was an overall trend towards clinical resolution among the challenge survivors, in all four groups.

**Fig. 5.**
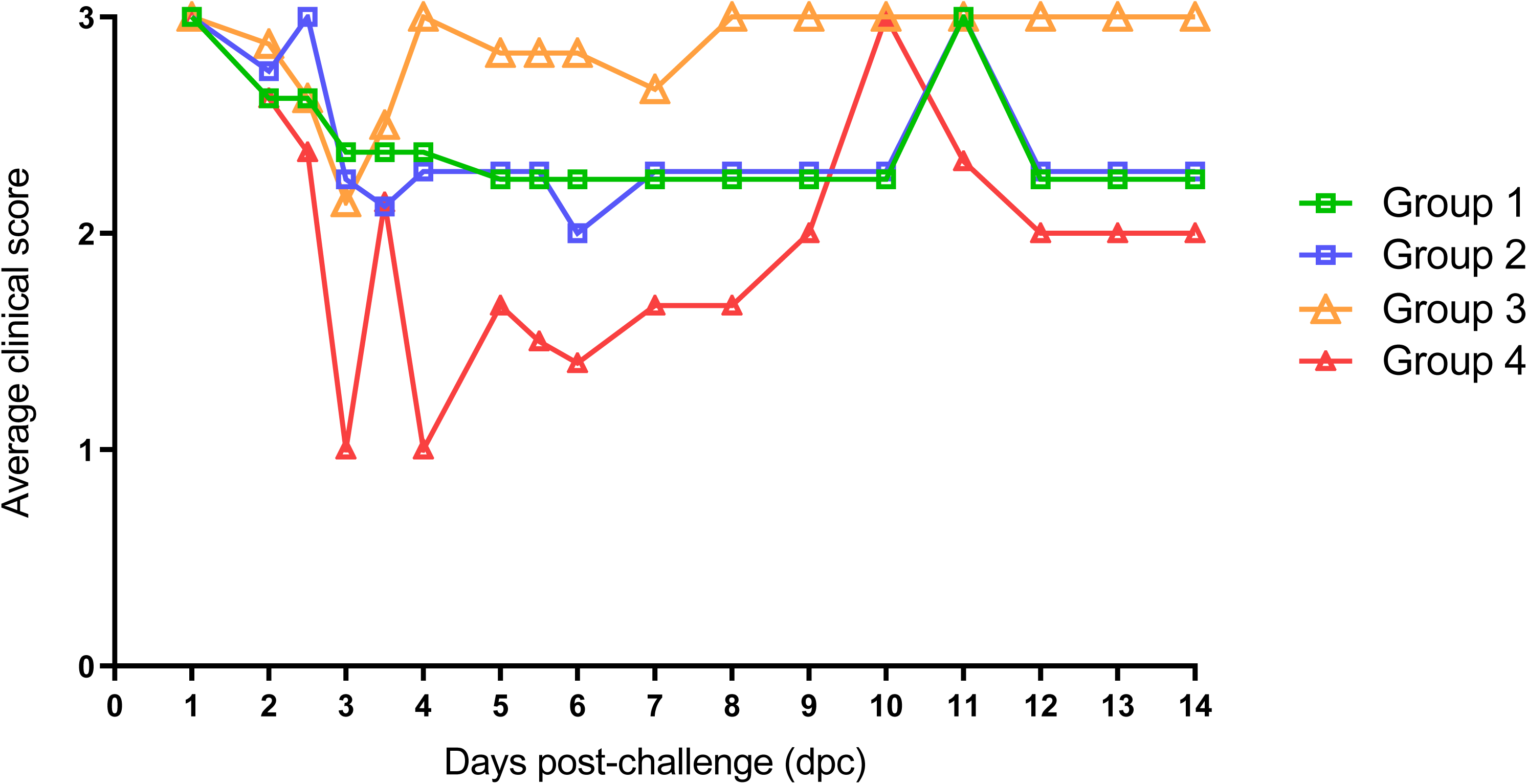
Scoring of clinical signs in the hens following H7N7-HPAIV challenge. The average daily clinical score for the surviving hens in each group is shown, with scoring frequency increased up to twice daily between 2-6 dpc, which spanned the period of the H7N7-HPAIV mortalities in all four challenged groups (Fig. 4). Individual hens were scored as experiencing either severe (requiring euthanasia, 0), moderate (1), mild (2) or no obvious clinical signs (3). Please refer to Methods for a summary of the observed signs.

### 3.4 H7N7-HPAIV pathogenesis

Following H7N7-HPAIV challenge, two apparently healthy hens from each group were culled at 2 dpc. Organ testing revealed H7N7-HPAIV dissemination by vRNA detection, although the proportions of infected organs varied among the four groups at 2 dpc (Figure 6a). The greatest systemic spread occurred in hens that succumbed to HPAIV infection in both the previously naïve hens (Group 4) and those previously inoculated with H7N7-DBCS (Group 2), with subsequent sampled deaths in these groups (2.4-3.4 dpc) also showing systemic dissemination. The least post-challenge systemic dissemination at 2 dpc occurred in two culled hens, previously inoculated with mall/SE/08 (Group 3): #3534 was apparently uninfected during the prior H7N7-LPAIV phase, and was devoid of any vRNA in all sampled organs at 2 dpc (Figure 6a). Hen #3533 experienced H7N7-LPAIV cloacal shedding only at 14 dpi (i.e. immediately prior to challenge), with low-level cloacal shedding observed at 1 dpc and its cull at 2 dpc. It was not possible to determine whether its shedding at 1-2 dpc was H7N7-LPAIV or H7N7-HPAIV, but organ testing at the 2 dpc cull revealed low-level vRNA in its duodenum, colon and bursa which inferred early-stage H7N7-HPAIV systemic dissemination. While #3534 and #3533 were apparently healthy at 2 dpc, the two subsequent Group 3 mortalities (#3540 and #3538) clearly experienced greater systemic H7N7-HPAIV dissemination (Figure 6a).

**Fig. 6.**
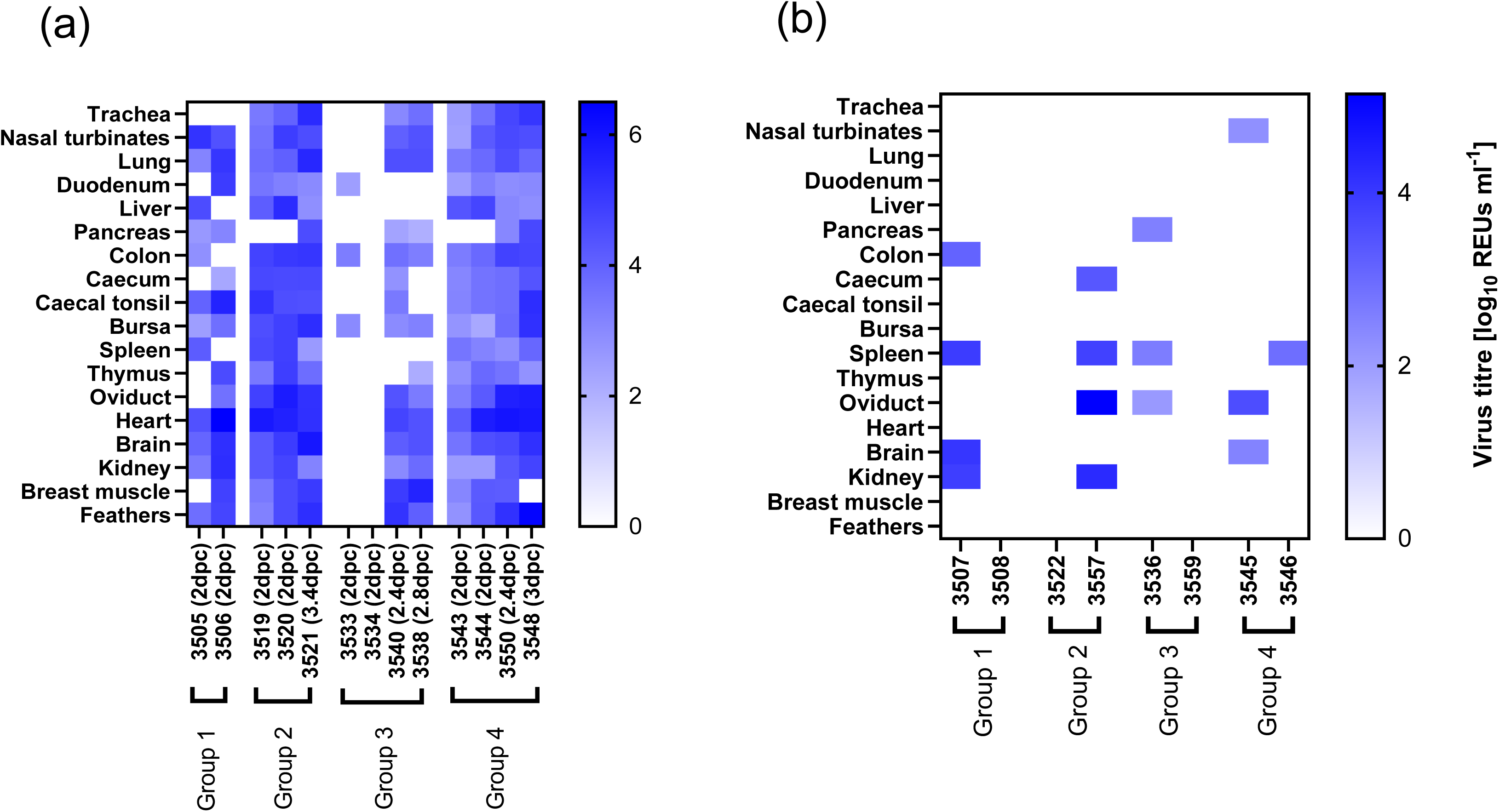
Viral RNA investigation of H7N7-HPAIV organ tropism. Following challenge of Groups 1-4. total RNA extracts from 18 organs per sampled hen were tested by M-gene RRT-PCR. Viral RNA levels indicated for (a) apparently healthy hens culled at 2 dpc in Groups 1-4 and any subsequent deaths which occurred in Groups 2-4; and for (b) hens in Groups 1-4 which survived until study-end (14 dpc) when they were culled. Viral RNA levels are shown as REUs, indicated by the scale adjacent to the heat-maps, with white indicating “No Ct” or REU values below Ct 36.

Among the 24 post-challenge survivors at study-end (14 dpc), two from each group were randomly culled for organ sampling. H7N7-HPAIV shedding had resolved or was resolving at this stage in all four groups, with vRNA detected in fewer tissues compared to earlier times post-challenge (Figure 6a). Organs from hens #3508, #3522 and #3559 (Groups 1-3 respectively) were devoid of detectable vRNA (Figure 6b). Interestingly, #3508 was the only surviving hen in the whole study which did not shed any H7N7-HPAIV following challenge, which underlined the protective effect of its prior infection and seroconversion following earlier H7N7-SBCS inoculation. Hens #3522 and #3559 shed H7N7-HPAIV following challenge, albeit the latter shed at only sub-threshold levels. Two Group 4 survivors also revealed fewer H7N7-HPAIV affected organs at 14 dpc, compared to the frequent tissue tropism in four Group 4 hens sampled earlier (Figure 6).

Virus-specific IHC investigation of tissues from hens culled at 2 dpc from the four groups affirmed H7N7-HPAIV tissue tropism derived from vRNA detection, albeit IHC staining sensitivity was lower than vRNA detection (Figures 6a and 7a). Similar patterns of extensive viral antigen detection were observed among mortalities between 2.4-3.4 dpc (Figure 7b), with example IHC images included (Figure S1). Reduced viral antigen detection in tissues at 14 dpc again suggested clearance (or ongoing clearance) of H7N7-HPAIV infection (Figure 7c). No histopathological changes were apparent at 2 dpc in any of the four groups, but subsequent mortalities in Groups 2-4 frequently included splenic and pancreatic necrosis, plus tracheitis. Other than multifocal areas of malacia which were confirmed by histopathology, no obvious gross lesions were observed at post-mortem in mortalities among Groups 2-4. At 14 dpc, the four Group 2 and Group 4 survivors revealed residual lesions, including nephritis, peritonitis and air saculitis.

**Fig. 7.**
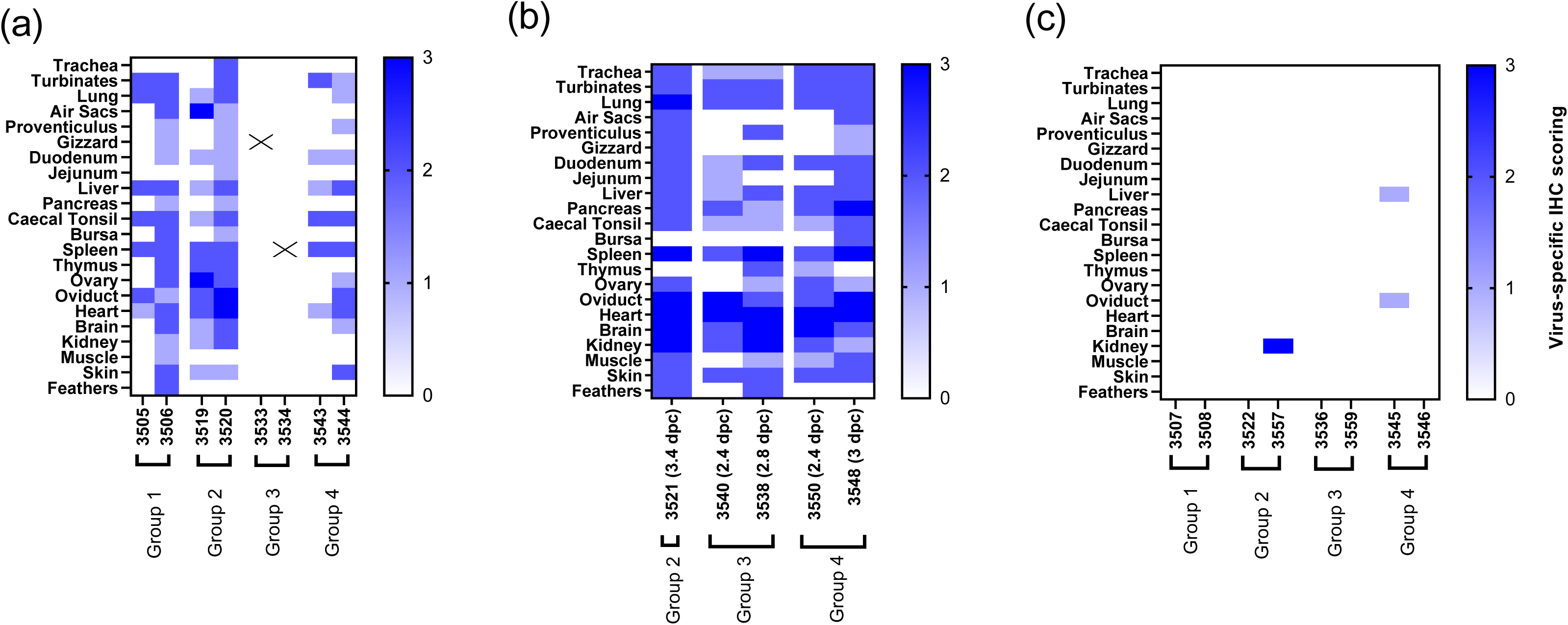
Virus-specific IHC investigation of H7N7-HPAIV organ tropism. Following challenge of Groups 1-4, 22 organs per sampled hen were investigated. The extent of virus-specific staining is presented for the tissues obtained from (a) two hens from each group which were pre-planned for cull at 2 dpc; (b) the subsequent deaths which occurred in Groups 2-4 on the indicated days post-challenge; and (c) two hens from each group which survived to study-end at 14 dpc. The semi-quantitative assessment of virus-specific IHC staining was scored as 0 (no staining), 1 (occasional observation of immunolabelled cells), 2 (moderate observation of immunolabelled cells) and 3 (numerous immunolabelled cells observed). “X” indicates that IHC staining was not done.

Interestingly, in hens #3508 and #3559 from Groups 1 and 3 respectively, post-mortem examination at 14 dpc revealed ongoing ovulation. Hen #3508 was the single instance of clinical protection accompanied by absence of HPAIV infection conferred by earlier H7N7-SBCS infection; hence it successfully resisted H7N7-HPAIV challenge. Hen #3559 had also developed H7-HI reactivity by the end of the H7N7-LPAIV phase, with subsequent H7N7-HPAIV shedding being limited to sub-threshold levels, reflecting ameliorated infection following H7N7-HPAIV challenge. The oviducts of both these hens were devoid of H7N7-HPAIV at 14 dpc, but H7N7-HPAIV was detected in the oviducts of other sampled hens following challenge (Figures 6, 7 and S1).

### 3.5 Viral environmental contamination

Drinking water and mixed straw/faeces samples were collected daily. During the H7N7-LPAIV phase, no environmental contamination was detected in Groups 1 and 2 (Figure 8). However, in the Group 3 housing, vRNA was detected in the drinking water at 6 dpi and straw/faeces at 12 dpi (Figure 8). Both instances of water contamination were preceded by low-level shedding in Group 3, which was more frequent than in Groups 1 and 2 (Figure 2).

**Fig. 8.**
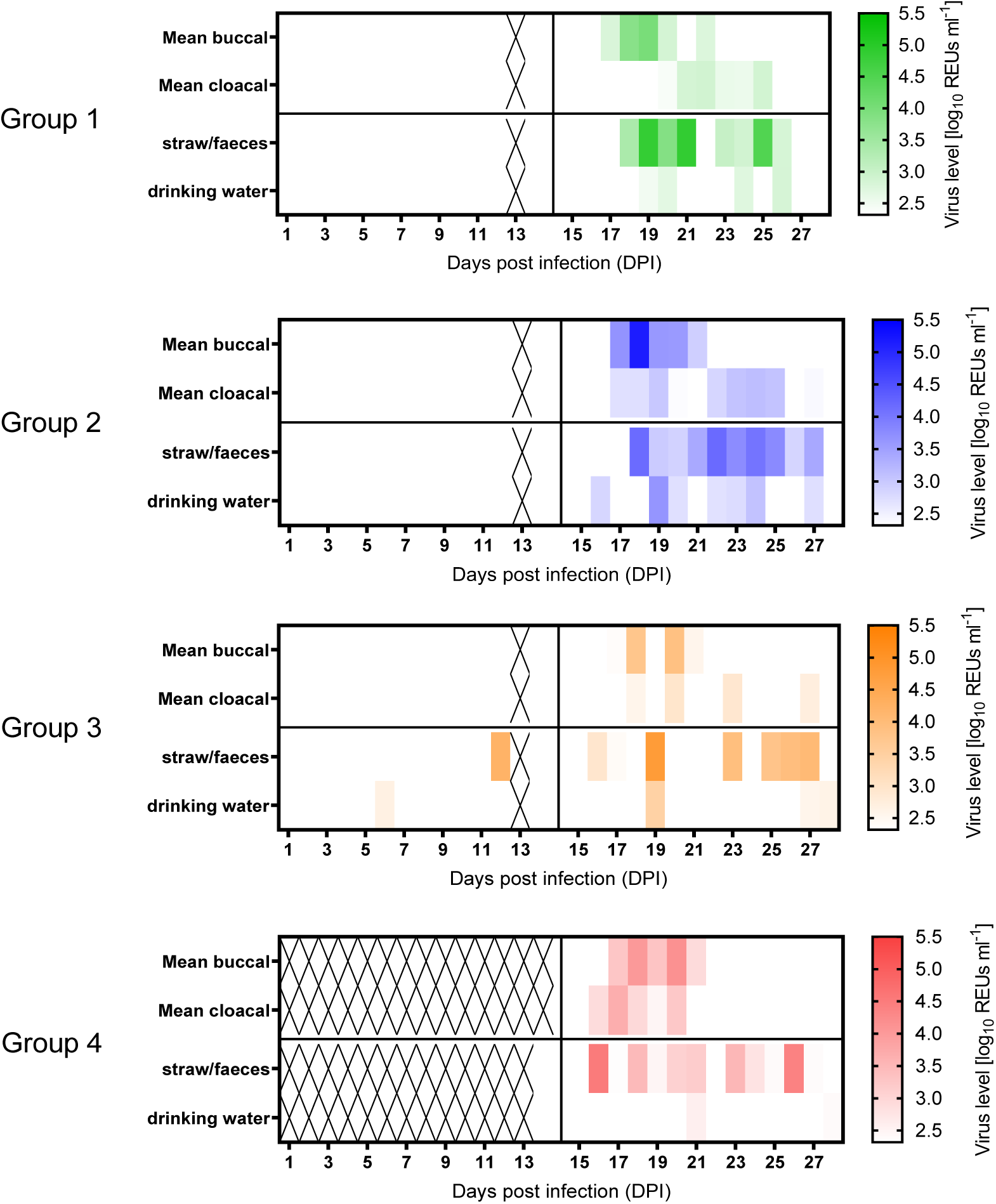
Viral contamination in the environment during the initial H7N7-LPAIV and subsequent H7N7-HPAIV phases. The four panels display the mean shedding (derived from Fig. 2) for the four corresponding Groups, together with vRNA levels detected by M-gene RRT-PCR in the straw/faeces and drinking water. The vertical line at 14 dpi indicates the H7N7-HPAIV challenge timepoint. Viral RNA levels are shown as REUs, indicated by the scale adjacent to the heat-maps, with white indicating “No Ct” or REU values below Ct 36. Both types of environmental samples were collected daily from each group’s housing. Drinking water was replaced daily, immediately after specimen collection, hence any H7N7 contamination in the water represents deposition during the preceding 24 hours, but the straw/faeces may have been contaminated at any time since the onset of shedding. “X” indicates that environmental samples were not collected just before the end of the H7N7-LPAIV phase at 13 dpi, while Group 4 sampling focused on the H7N7-HPAIV challenge phase, onwards from 14 dpi.

Following H7N7-HPAIV challenge, vRNA was detected in both environmental samples in all groups, on multiple days, which associated with ongoing shedding (Figure 8). Drinking water was refreshed daily, so any water contamination represented viral deposition during the preceding 24 hours. H7N7-HPAIV environmental contamination occurred less frequently in Groups 3 and 4 which experienced the higher mortalities following challenge. The greater reduction in the number of live (shedding) hens may have contributed to the less frequent H7N7-HPAIV water contamination in these two groups, but vRNA was detected in straw/faeces which may have been contaminated at any time since shedding onset.

## 4. Discussion

Although the importance of the HA CS mutation is long-known, there remain major knowledge gaps concerning HPAIV genesis (de Bruin et al., 2022), including the circumstances under which the switch to a MBCS occurs. Evidence garnered from an H7N7-HPAIV layer outbreak (UK 2008) identified a likely sequence of events, starting with prior H7N7-LPAIV incursion and spread which commenced approximately 2 weeks earlier (Seekings et al., 2018). These observations informed our novel study design, including initial inoculation of point-of-lay hens with potential H7N7-LPAIV precursors, before H7N7-HPAIV challenge at 14 dpi with the outbreak isolate, to reproduce the key HPAIV emergence event. Two H7N7-LPAIVs were produced by RG (Seekings et al., 2020), altering the outbreak HPAIV CS to the classical SBCS and the rare DBCS identified during the HPAIV outbreak, for initial inoculation of Groups 1 and 2, respectively. The third H7N7-LPAIV inoculation (Group 3) used a temporally related but genetically distinct European wild mallard isolate (mall/SE/08) with high similarity in the HA gene only, which included the DBCS (Metreveli et al., 2010; Seekings et al., 2018) Challenge outcomes revealed H7N7-HPAIV shedding in all four groups, but with varying clinical protection against HPAI mortality, acquired by the previously H7N7-LPAIV inoculated hens, with 100%, 87.5% and 75% survival in Groups 1-3, respectively. By comparison, the AIV-naïve challenge control hens (Group 4) experienced the lowest survival frequency of 37.5%.

Twenty-three of 24 (95.8%) hens in Groups 1-3 experienced H7N7-HPAIV shedding, indicating that prior experimental H7N7-LPAIV exposure did not prevent subsequent H7N7-HPAIV replication. These observations broadly concurred with the H7N7-HPAIV (2008) outbreak investigation, where prior exposure to the H7N7-LPAIV precursor (evidenced by frequent detection of H7-specific antibodies) did not totally protect hens against subsequent homologous H7N7-HPAIV infection and death (Seekings et al., 2018). H7-specific HI titres at the pre- and post-challenge stages revealed an overall increased humoral response among challenge survivors, at 14 dpc, as a likely boosting effect following H7N7-HPAIV challenge. The H7N7-HPAIV infected and surviving hens in Groups 1-3 (20/24; 83.3%) would represent, in a commercial poultry setting, a source for potential onward H7N7-HPAIV spread.

Despite investigating 82 clinical and faecal specimens during the UK H7N7-HPAIV (2008) outbreak, the immediate H7N7-LPAIV precursor was not isolated. The LPAIV DBCS was identified by partial HA sequencing from litter-derived specimens, but no full-genome sequences were obtained (Seekings et al., 2018). The H7N7-DBCS RG virus shared the identified CS sequence, but all other genetic segments were from the H7N7-HPAIV isolated at the outbreak. The LPAIV phenotype of both the H7N7-SBCS and H7N7-DBCS RG viruses was previously confirmed (Seekings et al., 2020), and our current in vivo study affirmed that MBCS removal from the HA gene ameliorated pathogenicity. However, by including sub-threshold shedding after H7N7-LPAIV inoculation, only one (12.5%) and two (25%) of the hens in the H7N7-SBCS (Group 1) and H7N7-DBCS (Group 2) became H7N7-LPAIV infected, respectively. The remaining 13 hens in both groups remained H7N7-LPAIV uninfected (Table 1), based on the absence of any shedding and no H7-HI serological reactivity, by 14 dpi.

Inoculation with the native-origin mall/SE/08, resulted in five hens (62.5%, Group 3) becoming infected, with more frequent shedding observed during the H7N7-LPAIV phase compared to the two RG H7-LPAIVs (Figure 2). This observation raised the possibility that the two RG-derived viruses used to inoculate hens in Groups 1 and 2 may not have been optimally adapted for efficient in vivo replication. The RG rescue of these LPAIVs was based on modification of only the CS in the HA gene. The authentic outbreak precursor H7N7-LPAIV may have contained unknown polymorphisms within the internal viral genes, which may have been more advantageous for replication and spread in layers, prior to H7N7-HPAIV emergence at the premises.

Age may have reduced H7N7-LPAIV replication efficiency as other studies used younger (3 to 6-weeks-old) chickens for H5/H7 LPAIV inoculations at similar doses, causing more frequent shedding (Claes et al., 2013; Mundt et al., 2009; Pillai et al., 2010; Post et al., 2013). However, similar inoculations of 4-week-old chickens with other H7-LPAIVs resulted in a range of infection frequencies (Kim et al., 2012; Lee et al., 2018; Spackman et al., 2010). Housing conditions at the H7N7-HPAIV layer outbreaks differed from those required by the high welfare standards in the current in vivo study, with field conditions potentially influencing increased susceptibility to H7N7-LPAIV infection (Chowdhury et al., 2019), prior to H7N7-HPAIV outbreak emergence. Chicken breed and genetics may also affect susceptibility to LPAIV infection and shedding dynamics (Ruiz-Hernandez et al., 2016).

While no evidence for an H7N7-SBCS precursor was discovered during the UK (2008) H7N7-HPAIV outbreak (Seekings et al., 2018), it was used in the current study as a potential precursor to H7N7-DBCS emergence, whereby H7N7-SBCS may have been outcompeted and became undetectable at the outbreak premises. Alternatively, the outbreak precursor may have directly incurred as the H7N7-DBCS, as suggested by isolation of the temporally related DBCS-containing mall/SE/08, during the same year. Other than mall/SE/08 and the H7N7-DBCS precursor discovered during the UK (2008) outbreak, the only other known H7-AIV with an HA DBCS was the H7N7-LPAIV isolated at an Australian (Victoria) duck farm in 1976, which preceded a H7N7-HPAIV outbreak at a neighbouring chicken farm (Bashiruddin et al., 1992; Turner, 1976). Related H7N7-LPAIV (SBCS) and HPAIV strains were successfully isolated from neighbouring layer outbreaks during 2015 in Germany, but were separated by 45 days (Dietze et al., 2018).

All eight Group 1-3 hens considered as H7N7-LPAIV infected survived subsequent H7N7-HPAIV challenge. Survivors included #3508 as the only hen which became H7N7-LPAIV infected (shedding) in Group 1 (H7N7-SBCS), and was the only hen among Groups 1-3 which seroconverted to a positive H7 HI titre (1:16; Figure 3) by the end of the H7N7-LPAIV phase. This hen was the only instance of complete protection against H7N7-HPAIV challenge, in that it not only survived, but was the only hen among all challenge groups which did not shed any H7N7-HPAIV. It remained healthy to study-end at 14 dpc, when its tissues were also negative for vRNA.

The H7N7-HPAIV challenge survivors among the H7N7-LPAIV uninfected hens, included all seven (100%), five of six (83.3%) and one of three (33.3%), in Groups 1-3 respectively. Although considered as H7N7-LPAIV uninfected, these 13 survivors may have benefited from some protective effect following earlier H7N7-LPAIV exposure, which was proportionately highest in Groups 1 and 2, despite absence of earlier H7N7-LPAIV shedding and associated humoral responses. A protective mechanism in these two groups may be due to identical internal genes shared by H7N7-SBCS, H7N7-DBCS and the H7N7-HPAIV challenge strain. Internal AIV proteins, particularly the polymerases, may elicit cell-mediated immunity which protects against mortality following subsequent HPAIV challenge (Seo and Webster, 2001).

Group 4 hens which were previously AIV-naive experienced 37.5% survival following H7N7-HPAIV challenge, by 14 dpc. All eight were H7N7-HPAIV infected, but the three survivors demonstrated resolution of H7N7-HPAIV shedding and clinical signs, along with successful seroconversion. In general, HPAI causes high mortality in AIV-naïve chickens (Alexander and Brown, 2009). However, experimental investigation of naïve 6-weeks-old chickens with H7N7-HPAIV (Netherlands, 2003) resulted in survival of 30% contact-infected chickens, which similarly resolved shedding (van der Goot et al., 2005). Direct intranasal inoculation of 2-weeks-old chickens with the Australian (Victoria 1976) H7N7-HPAIV resulted in 0% survival, but 20% survived among infected co-housed contacts which also seroconverted (Alexander et al., 1978). However, another investigation of the same H7N7-HPAIV differed in the chickens being 14-weeks-old, and resulted in up to 100% survivors among four groups of intranasal-inoculated and contact chickens, which all shed virus (Westbury et al., 1979).

However, experimental investigation of naïve 6-weeks-old chickens with H7N7-HPAIV (Netherlands, 2003) resulted in survival of 30% contact-infected chickens, which similarly resolved shedding (van der Goot et al., 2005). Direct intranasal inoculation of 2-weeks-old chickens with the Australian (Victoria 1976) H7N7-HPAIV resulted in 0% survival, but 20% survived among infected co-housed contacts which also seroconverted (Alexander et al., 1978). However, another investigation of the same H7N7-HPAIV differed in the chickens being 14-weeks-old, and resulted in up to 100% survivors among four groups of intranasal-inoculated and contact chickens, which all shed virus (Westbury et al., 1979). The survivors again all seroconverted after disease resolution. Therefore, survival of at least some previously AIV-naïve chickens following experimental H7-HPAIV infection has been described, with an indication that age may have a role in ameliorating H7N7-HPAIV mortality, where the Group 4 hens in our study were challenged at 19-weeks-age. However, such HPAI outcomes are likely to be a product of multiple factors, including the specific viral strain, while chicken genetics and immunological competence may have a role in disease resistance. As noted for the LPAIV phase of our study, the controlled conditions required by high welfare standards differed from those at chicken outbreaks, where experimentally housed chickens may also be more resilient to HPAI disease progression. While a detailed investigation of the underlying mechanism of HPAI resolution in AIV-naïve chickens was beyond the scope of our current experiments, the observation in Group 4 of clinical improvement, reduction in shedding and seroconversion at 14 dpc suggested that non-fatal infection was being cleared.

Another investigation of H7N7-HPAIV emergence from a H7N7-LPAIV precursor used mixed in vivo co-infections of chickens, with different proportions of the H7N7-HPAIV and the H7N7-LPAIV precursor (Germany, 2015 (Graaf et al., 2018)). The authors modelled how H7N7-HPAIV may emerge within individual chickens which experienced concurrent infection with the precursor H7N7-LPAIV. It was shown that H7N7-LPAIV outcompeted replication of the H7N7-HPAIV, and the latter only emerged successfully (as detected via chicken shedding) when the HPAIV:LPAIV ratio was much higher. It was speculated that a successful switch with HPAIV emergence in individual chickens likely occurred very soon after precursor LPAIV incursion, to avoid its inhibition by high levels of competing LPAIV replication (Graaf et al., 2018). Early HPAIV emergence is therefore likely to be followed by successful spread at the flock level, where low levels of homologous LPAIV adaptive immunity would enable HPAIV flock dissemination. In the current study, although the three H7N7-LPAIVs in Groups 1-3 did not infect 100% of the layers with (at most) only low-level and largely sporadic H7N7-LPAIV shedding, the low level H7-specific humoral responses elicited during the H7N7-LPAIV phase may represent partial homologous immunity, which did not prevent subsequent H7N7-HPAIV shedding in the majority of challenged hens. While a higher H7N7-LPAIV inoculation dose may have resulted in more frequent viral shedding, the partial immunity elicited by the H7N7-LPAIVs in Groups 1-3 may therefore be a realistic approximation of what occurred at the European layer outbreaks (Bonfanti et al., 2014; Byrne et al., 2021; Dietze et al., 2018; Mulatti et al., 2017; Seekings et al., 2018), where the H7N7-HPAIV emerged successfully.

AIV environmental persistence has been long suspected as contributing to outbreaks within poultry premises and as a risk for onward spread (Swayne, 2008). During the H7N7-LPAIV phase, only the Group 3 environment had occasional evidence of viral contamination, while those of Groups 1 and 2 did not. This observation likely reflected the greater number of mall/SE/08 infected (shedding) hens in Group 3, compared to the weaker and less frequent H7N7-LPAIV shedding recorded in Groups 1 and 2 (H7N7-SBCS and H7N7-DBCS inoculated, respectively). In another study, environmental contamination also correlated with shedding from H7N9-LPAIV infected chickens and turkeys (James et al., 2024). We also showed that frequent H7N7-HPAIV environmental contamination followed H7N7-HPAIV shedding, in all four groups. The clear demonstration of H7N7 environmental contamination emphasised the importance of effective biosecurity and disinfection at poultry premises (Swayne, 2008), particularly in view of continuing outbreaks, caused by both H5 and H7 subtypes (FAO, 2025).

The circumstances under which HPAIVs emerge from their homologous LPAIV precursor remain a major knowledge gap in the understanding of changing AIV pathogenesis. We demonstrated how evidence obtained from poultry outbreaks can inform appropriate experimental approaches to provide insights into the underlying events, which occur at novel HPAIV outbreaks. In addition to the European H7N7-HPAIV outbreaks which informed this study, H7-HPAIV poultry outbreaks in the USA and Mexico since 2009 have been similarly associated with an H7-LPAIV precursor (Killian et al., 2016; Lee et al., 2017; Maurer-Stroh et al., 2013; Youk et al., 2020). Our study underlined the importance of sequential events, which followed from initial H7N7-LPAIV circulation in layers and resulted in H7N7-HPAIV outbreaks, within 2 weeks. We demonstrated that initial H7N7-LPAIV exposure did not prevent subsequent homologous H7N7-HPAIV shedding, following challenge, in the majority of hens. The pathway of H7-HPAIV emergence is crucial not only for the understanding of the LPAIV to HPAIV switch, but also to inform the management and control of such economically damaging poultry diseases.

## Acknowledgement

The authors would like to thank the Animal Sciences staff at APHA-Weybridge for the collection of samples and assisting with the in vivo experiment.

