## Supplementary material for "Infection of point-of-lay hens to assess the sequential events during H7N7 high-pathogenicity avian influenza emergence at a layer premises": Suppl Fig S1

### Slide 1
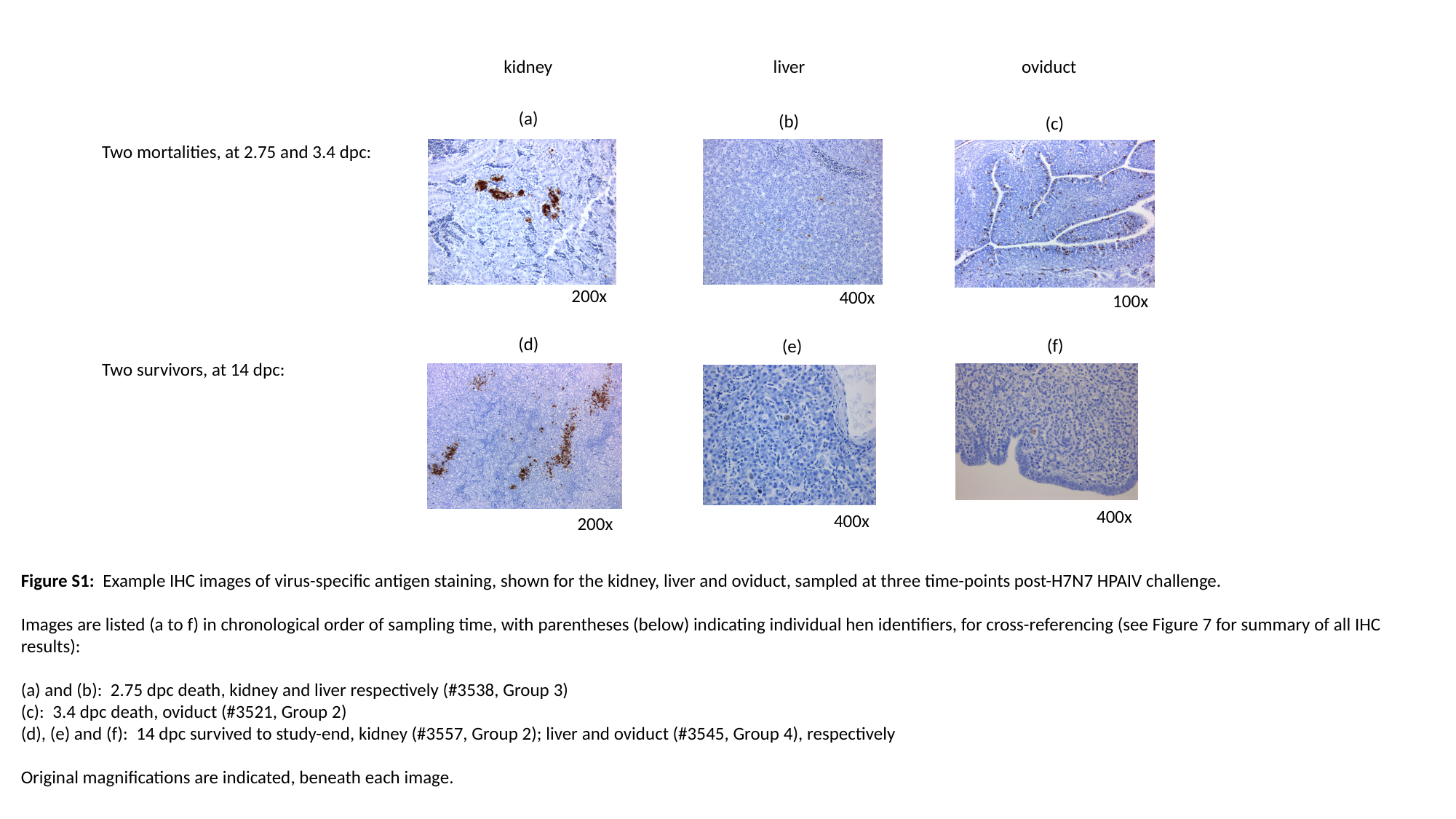

kidney liver oviduct
(a)
(b)
(c)
Two mortalities, at 2.75 and 3.4 dpc:
Two survivors, at 14 dpc:
200x
400x
100x
(d)
(f)
(e)
400x
400x
200x
Figure S1: Example IHC images of virus-specific antigen staining, shown for the kidney, liver and oviduct, sampled at three time-points post-H7N7 HPAIV challenge.
Images are listed (a to f) in chronological order of sampling time, with parentheses (below) indicating individual hen identifiers, for cross-referencing (see Figure 7 for summary of all IHC results):
(a) and (b): 2.75 dpc death, kidney and liver respectively (#3538, Group 3)
(c): 3.4 dpc death, oviduct (#3521, Group 2)
(d), (e) and (f): 14 dpc survived to study-end, kidney (#3557, Group 2); liver and oviduct (#3545, Group 4), respectively
Original magnifications are indicated, beneath each image.
